# Genetic Association of *CYP1A2* Gene Variant (rs762551) with Caffeine Induced-Hypertension Susceptibility and Cytochrome P450 1A2 Protein Analyses

**DOI:** 10.1101/2023.10.07.561327

**Authors:** Gulsher Amjad, Rashid Saif, Mehnaz Ghulam Hussain

## Abstract

High blood pressure is one of the most common illnesses affecting the Pakistani population due to, but not limited to, dietary habits, sedentary lifestyle and socio-economic aspects which have devastating effects on human health and general well-being. Caffeine is metabolized by CYP1A2 which results in lowered blood pressure while its retention in case of low metabolism may lead to hypertension. This phenomenon occurs because caffeine and its metabolites block A1 receptors in the kidney lowering the function of adenosine in blood pressure regulation. Current research aimed to demonstrate the genetic association of *CYP1A2* gene variant 15:74749576C>A (C allele lowering CYP1A2 activity) within caffeine induced hypertensive individuals of Pakistani origin using ARMS-PCR. This pilot scale study revealed that overall, 8%, 56% and 36% sampled population (n=50) is homozygous wild-type (C/C), heterozygous (C/A) and homozygous mutant (A/A) respectively. Similarly, alternative allele frequency is 0.28 and 0.48 in cases and controls. Chi-square (χ^2^) association test using PLINK data analysis toolset was applied which showed significant results of χ^2^(2, *N* = 50) = 4.244, *p* = 0.039. Hardy Weinberg Equilibrium analysis was also applied which establishes that sampled population is obeying the principle with *p-value* of 0.241. Moreover, odds-ratio depicts that the mutant allele is 0.42 times less prevalent in cases vs. controls. A few bioinformatics tools were employed e.g., ProtParam, PsiPred, PDB-RCSB, Motif finder, CTU-TMHMM-2.0, ScanProsite, GPS PAIL2.0, PRmePRed, NetOGlyc4.0, NetPhos3, SIFT analysis and STRING database in order to predict physicochemical properties, secondary structure, 3-dimensional structure, conserved motifs, transmembrane structure, post-translational modifications, protein variants impact on its function and protein-protein interactions respectively. The current endeavor attempted to provide the genetic architecture of the aforementioned variant in this case-control study in Pakistani individuals, which can pave paths of preventive medicine initiatives through genetic counsel of the masses.

## Introduction

Hypertension is referred to blood pressure higher than the global average range of <120/80 mmHg. According to National Diabetes Survey of Pakistan, prevalence of hypertension in urban and rural areas was 44.3% and 46.8% in 2020 respectively. Overall, age-adjusted weighted prevalence of hypertension was 46.2%, of which 24.9% were self-reported while 21.3% newly diagnosed [1]. Hypertension symptoms may include; headache, heart palpitations, nosebleeds, nausea, muscle and chest pains, which may have severe consequences such as strokes, heart attacks, kidney failures and overall poor routine of an individual. Pakistani population is also on the forefront of this ailment due to many reasons varying from environmental to genetic tendencies as well as casual and conservative behavior to cope with, rather than taking proper recommended drugs to cure this life-threatening peril. This study keeps in consideration the relationship of *CYP1A2* and caffeine intake as it is common among Pakistani inhabitants through to the consumption of black tea as well as inhaling of cigarette smoke.

The Cytochrome P450 1A2 (*CYP1A2*) gene was selected in the current study due to numerous factors like its impact on several pathways which are associated to hypertension and the fact that *CYP1A2* is expected to mediate 90% of the primary caffeine metabolism [2]. The CYP1A2 enzyme is involved in the metabolism of many drugs e.g., (clozapine, theophylline, tacrine and caffeine) as well as several other endogenous compounds e.g., (melatonin, estrone and estradiol). The CYP1A2 protein also plays a role in the bioactivation of various toxic compounds and procarcinogens e.g., (aflatoxin B1, polycyclic aromatic hydrocarbons and aromatic or heterocyclic amines) [3]. There are various CYP1A2 enzyme activity levels that have been linked to different SNPs in the *CYP1A2* gene. [4]. In normal conditions, Adenosine binds to A1 receptors in the kidney and promote Na^+^ and water reabsorption. However, in the case of high CYP1A2 activity, caffeine gets readily metabolized and both caffeine along with its metabolites antagonize the A1 receptor resulting in decreased Na^+^ and water reabsorption which could potentially lower blood pressure[5].

A plethora of variants have been reported in the literature which are associated to hypertension with significant *p*-values in different populations. Aforementioned *CYP1A2* gene is located on Chr.15 NC_000015.10 at 15q24.1 locus. Genomic position of the selected variant is g.74749576C>A according to genome assembly (GCF_000001405.40) with locus c.-9-154C>A, global minor allele frequency of (C) is 0.37021. This particular gene (transcript ID: NM_000761.5) has a total of 7 exons with encoded protein ID NP_000752.2 The subject protein has 516 amino acids (aa) of which 147^th^ aa is affected by the selected variant along with the splice distance of 154 nucleotides downstream of the variant[6]. According to ClinVar it is likely benign in nature[7]. Furthermore, the dbSNP ID is rs765521 which is on intron 1 at 679 of total 832nt. Individuals with rs762551 C allele have been demonstrated to have decreased CYP1A2 activity [8], which may make them more vulnerable to high blood pressure and hypertension.

Here the subject variant is being genotyped using ARMS-PCR followed by *HWE*, χ^2^ and OR statistics for determining the association and correlation of this variant with hypertensive individuals from Pakistan.

## Materials and Methods

### Sample collection and DNA extraction

A total of 50 blood samples were genotyped, including samples from patients suffering from hypertension (n=25) and control samples (n=25), to look for a connection between the *CYP1A2* locus and hypertension within Pakistani Population. The blood samples of various gender and age groups were collected in K3-EDTA vacutainers and stored at 4°C until further use. TIANAMP Genomic DNA Kit (TIANGEN Biotech Beijing Co., Ltd.) was used to extract DNA following manufacturer’s instructions.

### ARMS-PCR primer designing

Primers were designed for *CYP1A2*, drawing up its properties from NetPrimer software (https://www.premierbiosoft.com/netprimer/). Both wild and mutant type variants were amplified using ARMS-PCR with the mutant type containing a mismatch 4 bases away from the 3’ of the gene. Moreover, internal control (IC) primers are used to ensure PCR accuracy (Table 1).

**Table 1:**
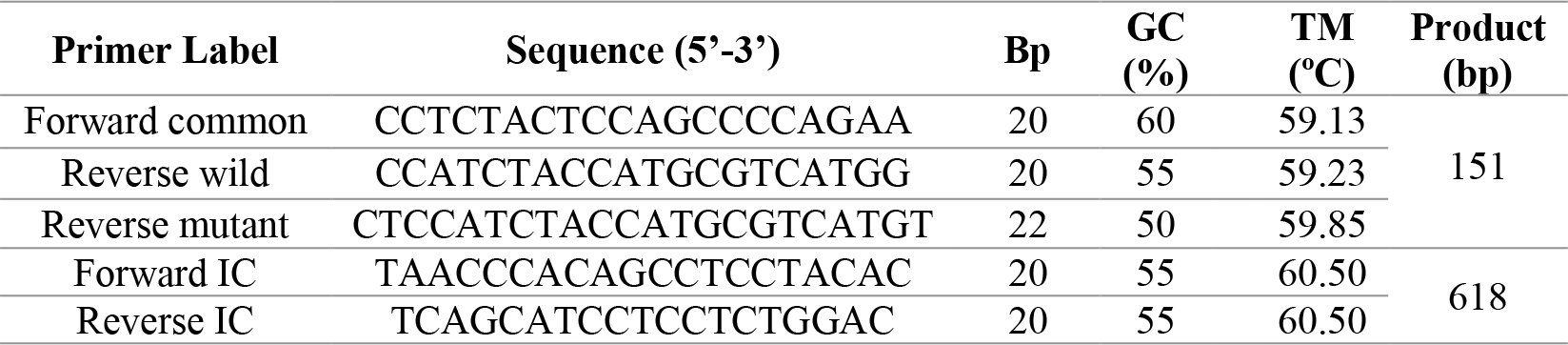
ARMS-PCR primer sequences and their properties.

### PCR Amplification

Each sample (wild type and mutant variants) was amplified using SimpliAmp thermocycler (Applied Biosystems). For each sample, two PCR reactions were performed, each containing Forward Common primer with reverse wild and mutant type primers separately. Simultaneously, forward and reverse IC primers were also used to amplify internal control regions. A total of 12μL of reaction mixture was prepared consisting 2μL of 50ng/μL genomic DNA, 10mM of each primer, 0.015IU/μL of *Taq* polymerase, 2.5mM MgCl_2_, 2.5mM dNTPs, 1x *Taq* buffer and PCR-grade water. The PCR protocol started with 5-minute initial denaturation at 94°C followed by 25 cycles of denaturation at 94°C for 45 seconds, annealing at 59°C for 30 seconds, extension at 72°C for 30 seconds with the final extension at 72°C for 10 minutes and storage at 4°C (Figure 1).

**Figure 1:**
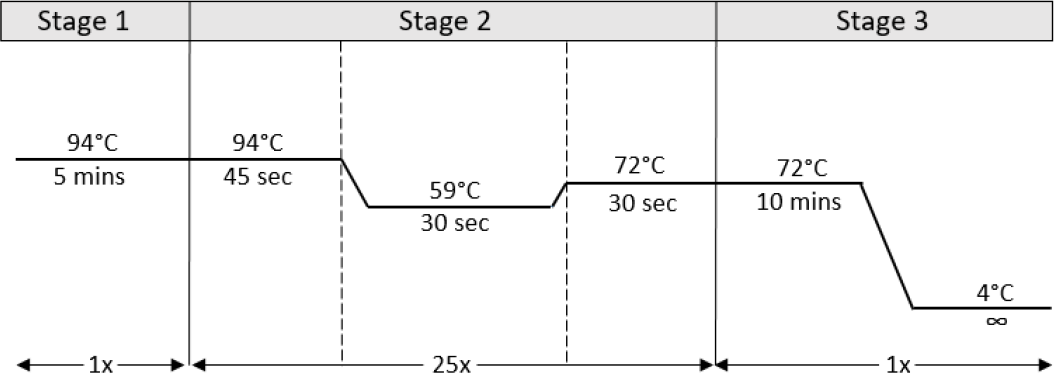
Thermal cyclic conditions of ARMS-PCR

### Statistical analysis

PLINK toolset was used to calculate the observed and expected genotyping frequencies, taking into account *HWE* by following *p*2 + 2*pq*+ *q*2 = 1 equation and Chi-square analysis test using 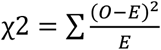 to confirm the association between subject variant 74749576C>A and CYP1A2 activity in the samples. Moreover, *p-* value and OR was also calculated together with allelic frequencies.

### Bioinformatics analysis

Different bioinformatics tools/databases were used to observe the various properties of wild-type CYP1A2 protein in the current study under an *in-silico* approach. The CYP1A2 protein transcript was acquired from NCBI. The tools were not used on mutant type protein to avoid redundant analysis because the mutation was synonymous in nature.

### Physicochemical properties

The physicochemical properties, theoretical PI, amino acids composition, atomic composition, molecular weight, instability aliphatic index, GRAVY, extinction coefficient and estimated half-life in the wild-type CYP1A2 protein sequences were figured out by uploading the FASTA sequence into “ProtParam” tool (http://web.expasy.org/protparam/) after which, the results were concluded.

### Secondary structure predictions

“PsiPred” (http://bioinf.cs.ucl.ac.uk/psipred/) was used to predict the secondary structure (alpha helices, beta sheets and coils) of the wild-type protein by uploading the FASTA sequence of the protein with default parameter. Visual Molecular Dynamics (VMD) was used to view the secondary structure of the subject protein.

### 3D protein structure

“PDB RCSB” (https://www.rcsb.org/) tool was used to visualize and analyze the protein’s 3D structure. The FASTA sequence was uploaded onto the database and a 3D image was generated.

### Protein conserved motifs prediction

The conserved sequences in the wild-type protein were gauged using the “Motif finder” tool. First, the protein sequence of the protein was searched and then the option for “motifs” was selected.

### Transmembrane structures estimation

To predict the transmembrane helices, intracellular and extracellular regions of the wild-type protein, the FASTA sequence of the CYP1A2 protein was uploaded onto “CTU-TMHMM” server v.2 (https://services.healthtech.dtu.dk/service.php?TMHMM-2.0)

### Post-translational modifications assessments

To check for various post-translational modifications in the wild-type CYP1A2 protein various tools were utilized such as “ScanProsite” (http://prosite.expasy.org/scanprosite/) for phosphorylation, “GPS_PAIL2.0” (http://bdmpail.biocuckoo.org/prediction.php) for acetylation, “PRmePRed” (http://bioinfo.icgeb.res.in/PRmePRed/#) for methylation and “NetOGlyc-4.0” (https://services.healthtech.dtu.dk/service.php?NetNGlyc-1.0) for glycosylation

### Assessment of impact of variants on protein function

Sorting Intolerant from Tolerant (SIFT) tool was used to measure the effect of non-synonymous SNPs and the pathogenicity of gene variant 74749576C>A of wild-type CYP1A2 protein. A mutation is deemed deleterious if the score is <0.05

### Protein-protein interactions illustrations

The STRING database/tool was used to figure out the protein-protein interface. The protein sequence was inserted into the tool and the results generated were used to analyze individual protein reactions.

## Results

A total of 50 blood samples were genotyped, (cases = 25, controls = 25) and after experimental analysis, it was concluded that there was 01 homozygous wild-type (C/C), 12 heterozygous (C/A), and 12 homozygous mutants (A/A) in cases (Figure 2 a). Whereas there are 03 homozygous wild-type, 16 heterozygous, and 06 homozygous mutants in the control cohort (Figure 2 b). Therefore, the overall genotypic frequency came out to be 0.08 (8%) for homozygous wild-type, 0.56 (56%) for heterozygous and 0.36 (36%) for homozygous mutant. Afterwards, Chi-square analysis was conducted to confirm that the population obeys *HWE* with the following *p-value* = 0.039.

**Figure 2:**
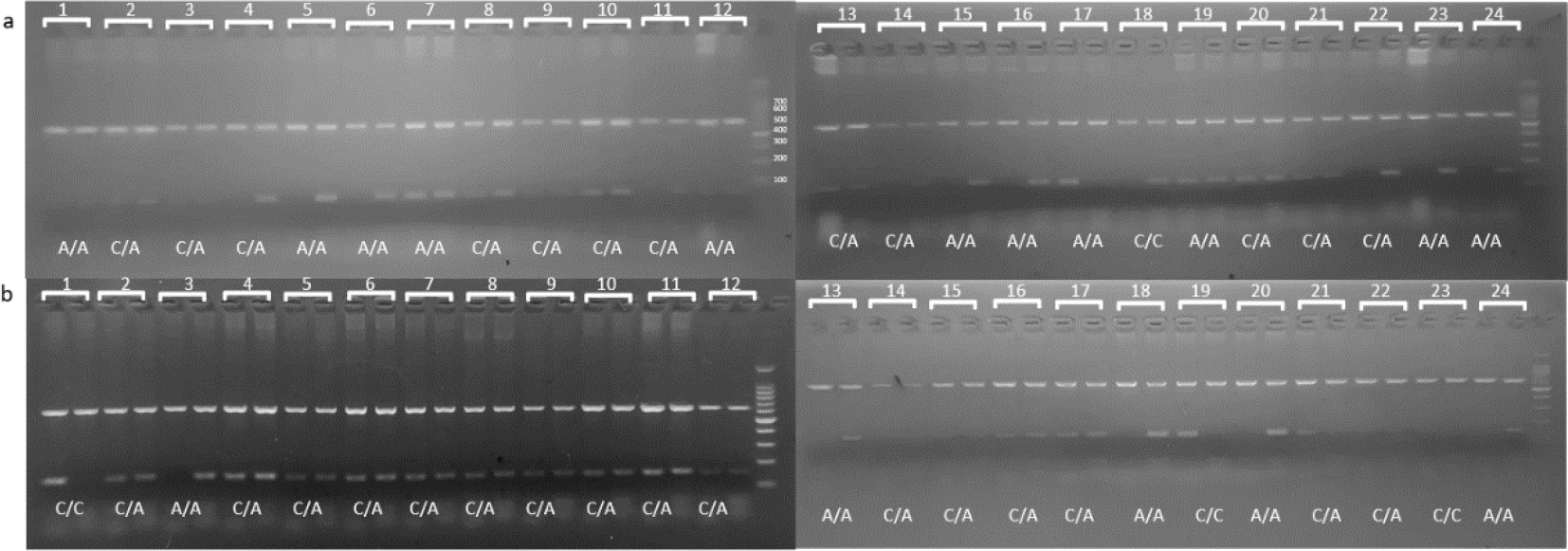
ARMS-PCR products of selected variant in 50 samples, a) cases, b) controls. Last lane of each gel contains 100 bp DNA standard.

### Statistical association analysis

Moreover, PLINK data toolset was used which showed that the alternative allele frequency is 0.28 (28%) and 0.48 (48%) in cases and control cohorts respectively. χ^2^ value of 4.244 and *p-value* of 0.039 shows significant genetic association between the selected variant with increased blood pressure as *p* < 0.05. Similarly, odds-ratio (OR) of 0.42 was observed suggesting that the prevalence of mutants is almost 0.42 times higher in controls as compared to cases (Table 2).

**Table 2:**
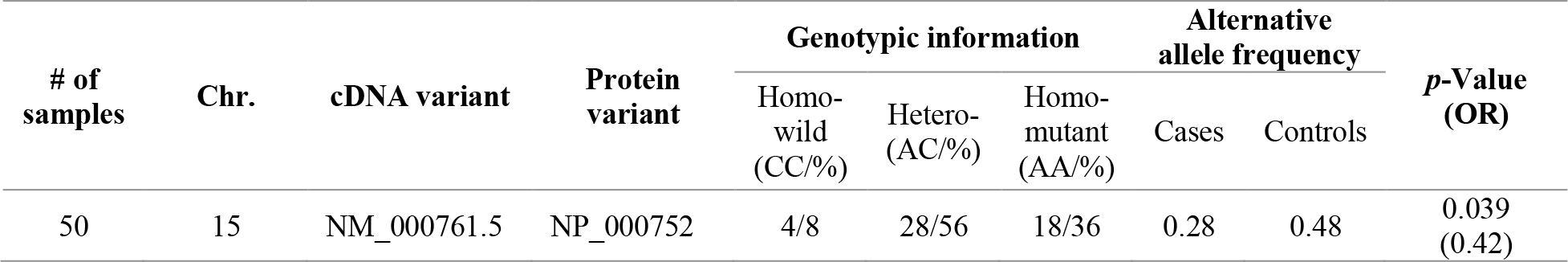
Association of subject variant with hypertension susceptibility in Pakistani population.

### CYP1A2 protein analysis using different bioinformatics tools

#### I. Physicochemical findings of CYP1A2 by ProtParam

Physicochemical parameters of CYP1A2 were analyzed by ProtParam tool and it was observed that the protein consists of 516 amino acids with theoretical PI 9.18 and 5840.7.47 Daltons molecular weight. The calculated instability index was 40.63 which predicted unstable protein with an estimated half-life of 30hrs. Additionally the extinction coefficient of 64775 was observed forming cysteine residues and 64400 assuming all pairs of cysteine are reduced. The protein has molecular formula C_2652_H_4144_N_718_O_763_S_17_ with positively charged Arg + Lys residues value of 59 and negatively charged Asp + Glu value of 49. Grand average of hydropathicity (GRAVY) was -0.153 (Table 3).

**Table 3:** Physicochemical properties of wild-type CYP1A2 protein.

| Physicochemical Properties | (Wild-type) |
| --- | --- |
| Amino acids | 516Aa |
| Molecular weight | 5840.7.47 |
| Theoretical PI | 9.18 |
| Molecular formula | C<sub>2652</sub>H<sub>4144</sub>N<sub>718</sub>O<sub>763</sub>S<sub>17</sub> |
| Number of atoms | 8267 |
| Negative charged residues (Asp + Glu) | 49 |
| Positive charged residues (Arg + Lys) | 59 |
| Extinction coefficient | 64775(assuming all pairs of Cys form cysteine)<br>64400(assuming all pairs of Cys are reduced) |
| Estimated half life | 30hrs |
| Instability index | 40.63 |
| Aliphatic index | 90.31 |
| GRAVY | -0.153 |

A detailed analysis of the protein sequence was done which showed that amino acid Leucine (Leu) was present in the highest amount, making up 12% of the protein. Valine (Val), Alanine (Ala), and Arginine (Arg), made up 6% of the protein individually while the other amino acids amounted to <6% (Figure 3).

**Figure 3:**
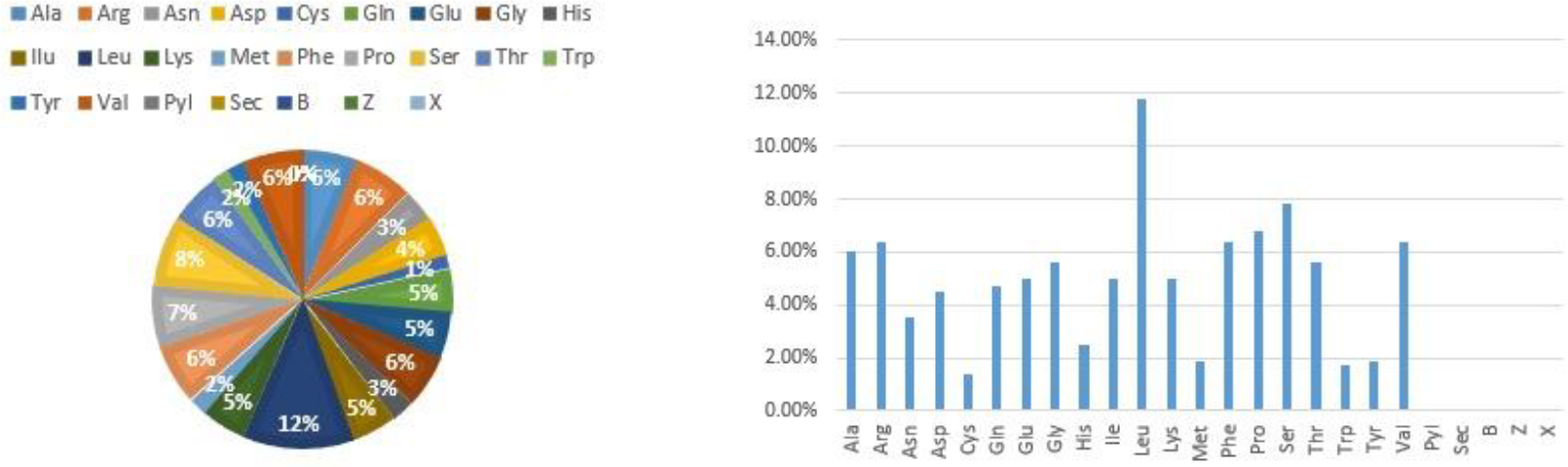
protein sequence constitution

#### II. Secondary structure predictions of CYP1A2

PsiPred tool predicted the secondary structure of CYP1A2 protein, the pink colored amino acids used for helices, yellow color for beta strands, whereas light grey for coils (Figure 4).

**Figure 4:**
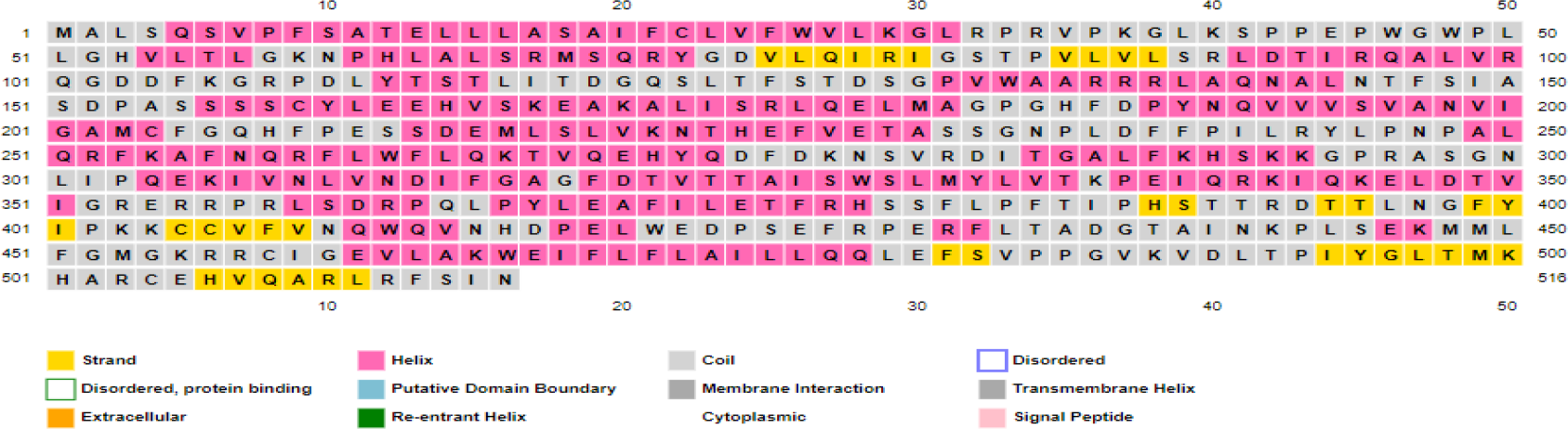
Amino acids involved in formation of alpha helices, beta strands and coils of the subject protein

#### III. 3D structure prediction of CYP1A2

The 3D crystal structure of human microsomal P450 1A2 in complex with alpha-naphthoflavone retrieved through X-RAY diffraction was extracted from PDB database. The structure illustrates alpha helices, beta strands and random coils which can be visualized in (Figure 5). To retrieve the pure structure of the protein, the attached group can be removed for downstream analysis i.e. molecular docking and dynamic simulation studies.

**Figure 5:**
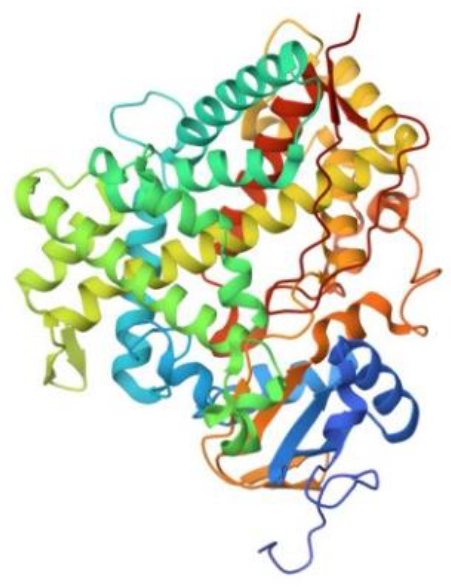
3D structure of human microsomal P450 1A2 in complex with alpha-naphthoflavone

#### IV. Protein Motifs predictions using Motif finder

Only one motif in CYP1A2 with pfam ID (P450) was predicted by Motif finder tool with the threshold E-value 42.492(6.8e-104) (Table 4). Motifs are short conserved sequence patterns which are associated with distinct structural properties and can function independently. They are particularly useful in protein classification and functional annotation.

**Table 4:** Motif prediction of CYP1A2 protein.

| pfam ID | Independent E-value (position) | Description |
| --- | --- | --- |
| P450 | 42..492(6.8e-104) | PF00067, Cytochrome P450 |

Furthermore, ScanProsite tool also depicted the graphical position of this motif as well as observing one hit in amino acid sequence FGMGRRCICEV. No disulfide bridge, active site or any other site was shown (Figure 6).

**Figure 6:**
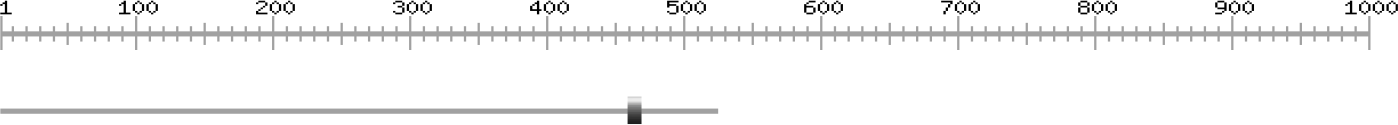
Post-translational modifications prediction of CYP1A2 protein

#### V. Transmembrane helices (TMHs) finding

According to TMHMM tool there are transmembrane regions which are shown in purple on the exterior side of the cell membrane, shown as an orange horizontal line, at the start of the sequence and few transmembrane regions inside the membrane, shown as a blue horizontal line, later on in the sequence (Figure 7). Transmembrane helices are important for transducing signals and providing a channel for ions through the lipid membrane.

**Figure 7:**
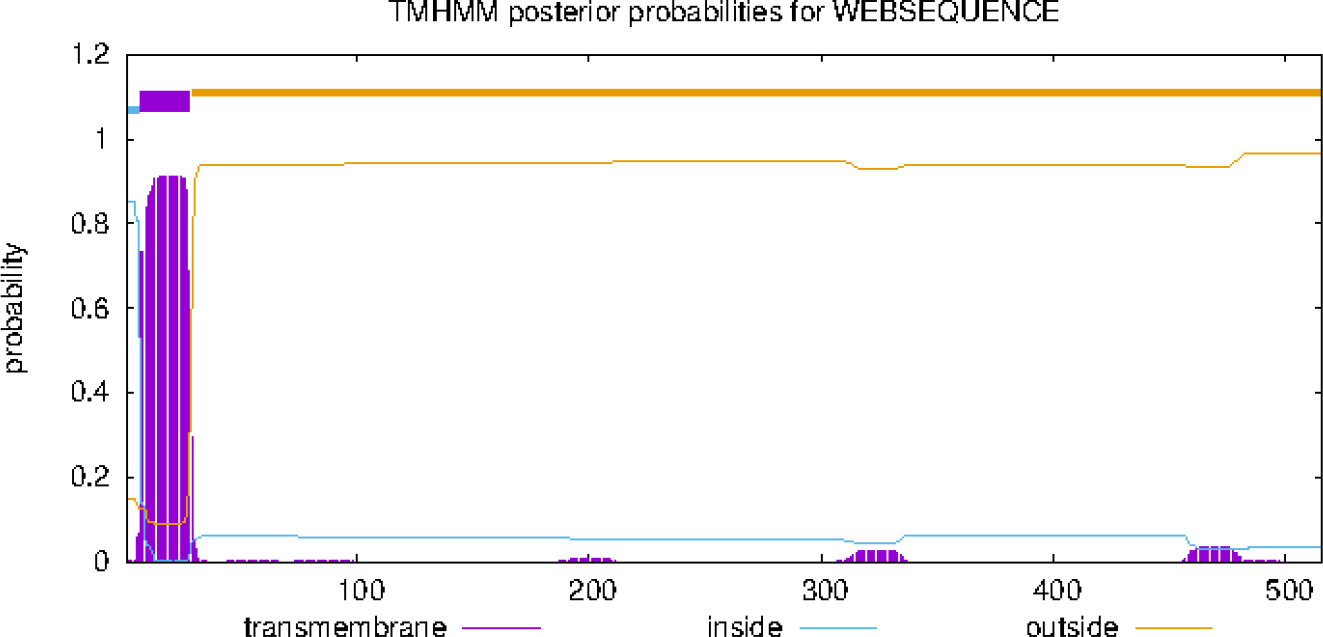
Transmembrane helices in CYP1A2 protein

#### VI. Post-translational modifications findings using ScanProsite

##### a. Glycosylation predictions of CYP1A2 using NetOGlyc-4.0

The graphical analysis of CYP1A2 by NETOGLYC-4.0 tool displayed the glycosylation on carbohydrates binding to target protein. The X-axis shows position of sequence while Y-axis shows N-glycosylation potential and purple line indicates the threshold value of 0.5 specifying the greater possibility of glycosylation if the threshold value is above 0.5. The yellow and black lines, if present, predict the potential and additional thresholds in between 0 and 1 respectively (Figure 8).

**Figure 8:**
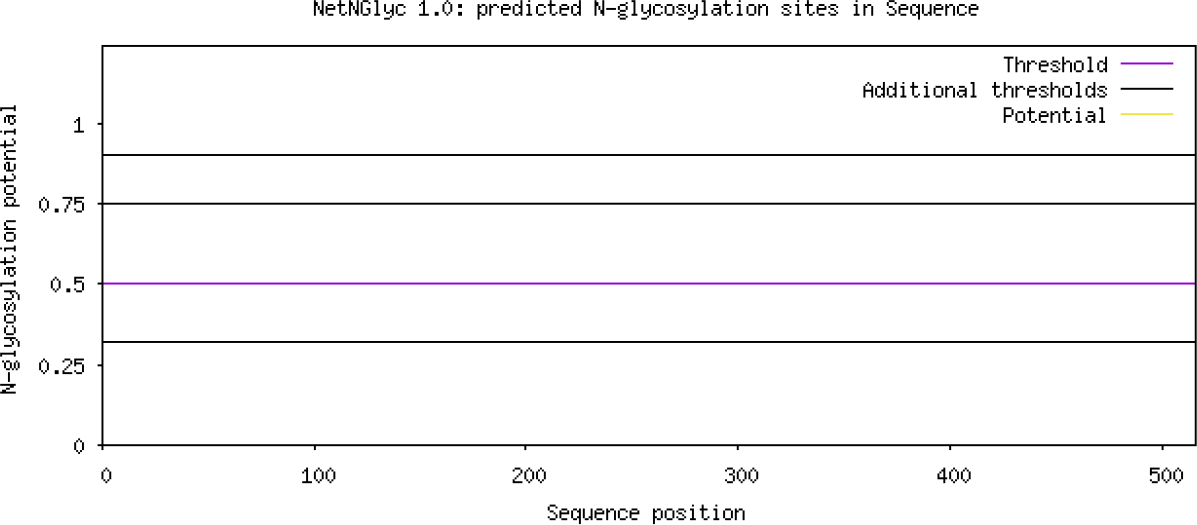
Glycosylation sites for CYP1A2 protein

##### b. R and K residues of methylation by PRmePRed

All the R and K residues of methylation of CYP1A2 protein were analyzed and predicted by PRmePRed tool at various positions because experimental determination of Arginine methylation sites is a time consuming and costly process. Table 5 contains different peptide sequences and prediction scores at various positions along the protein.

**Table 5:** Methylation sites of CYP1A2 protein.

| Position | Peptide | Prediction score |
| --- | --- | --- |
| 32 | LVFWVLKGLRPRGPKGLKS | 0.512985 |
| 34 | FWVLKGLRPRVPKGLKSP | 0.814597 |
| 67 | GKNPHLALSRMSQRYGDVL | 0.551364 |
| 79 | QRYGDVLQIRIGSTPVPVL | 0.547554 |
| 90 | GSTPVLVLSRLDTIRQALV | 0.562833 |
| 100 | LDTIRQALVRQGDDFKGRP | 0.697776 |
| 108 | VRQGDDFKGRPDLYTSTLI | 0.616363 |
| 136 | TDSPVWAARRRLAQNALN | 0.764639 |
| 137 | DSGPVWAARRRLAQNALNT | 0.545082 |
| 175 | SKEAKALISRLQELMAGPG | 0.588133 |
| 296 | LFKHSSKKGPRASGNLIPQE | 0.847605 |
| 353 | QKELDTVIGRERRPRLSDR | 0.900134 |
| 355 | ELDTVIGRERRPRLSDRPQL | 0.918855 |
| 356 | LDTVIGRERRPRLSDRPQL | 0.77547 |
| 358 | TVIGRERRPRLSDRPQLPY | 0.87469 |
| 362 | RERRPRLSDRPQLPYLEAF | 0.628961 |
| 457 | MMLFGMGKRRRCIGEVLAkW | 0.6628454 |
| 512 | RCEHVQARLRFSINXXXX | 0.653901 |

##### c. Acetylation of lysine (K) prediction in CYP1A2 by GPS PAIL 2.0

Acetylation by different histone acetyl transferases (HATs) is an important post translational modification which most notably plays an important role I n transcriptional regulation. In CYP1A2 protein the acetylation of Lysine (K) residue by a Lysine acetyl transferase (KAT) is at position K293 in peptide sequence ALFKHSKKGPRASGN with 100% KAT2A distribution sites score 1.667 and cutoff value 1.382 predicted by GPS PAIL 2.0 (Figure 9).

**Figure 9:**
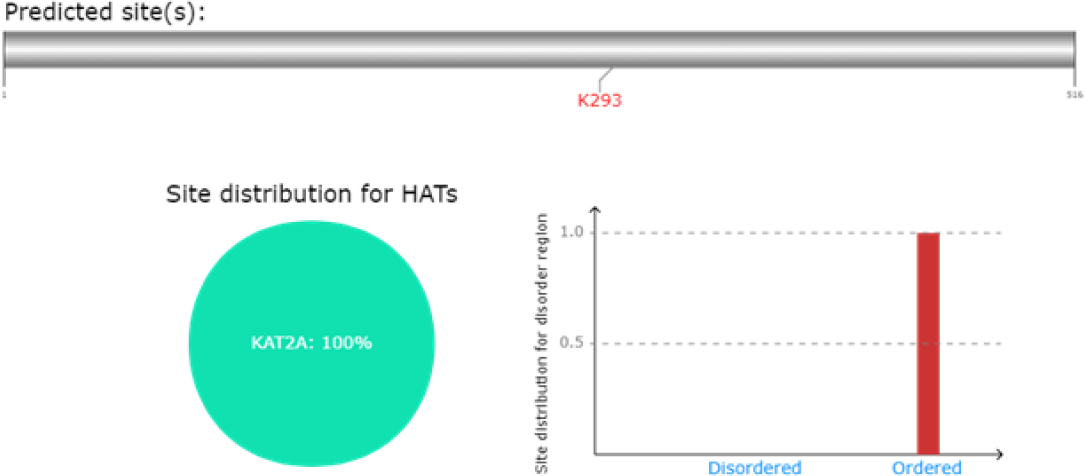
Acetylation of lysine(K) in CYP1A2 protein

##### d. Tyrosine, serine and threonine phosphorylation sites prediction by NetPhos 3.1

In wild-type CYP1A2 protein threonine, serine, and tyrosine phosphorylation sites were predicted by NetPhos 3.1 tool. In the graph, the pink line depicts the threshold limit of phosphorylation (0.5 for CYP1A2), red line shows serine phosphorylation, green line shows threonine phosphorylation and blue line shows tyrosine phosphorylation. Higher the threshold value more possibility of phosphorylation there is (Figure 10).

**Figure 10:**
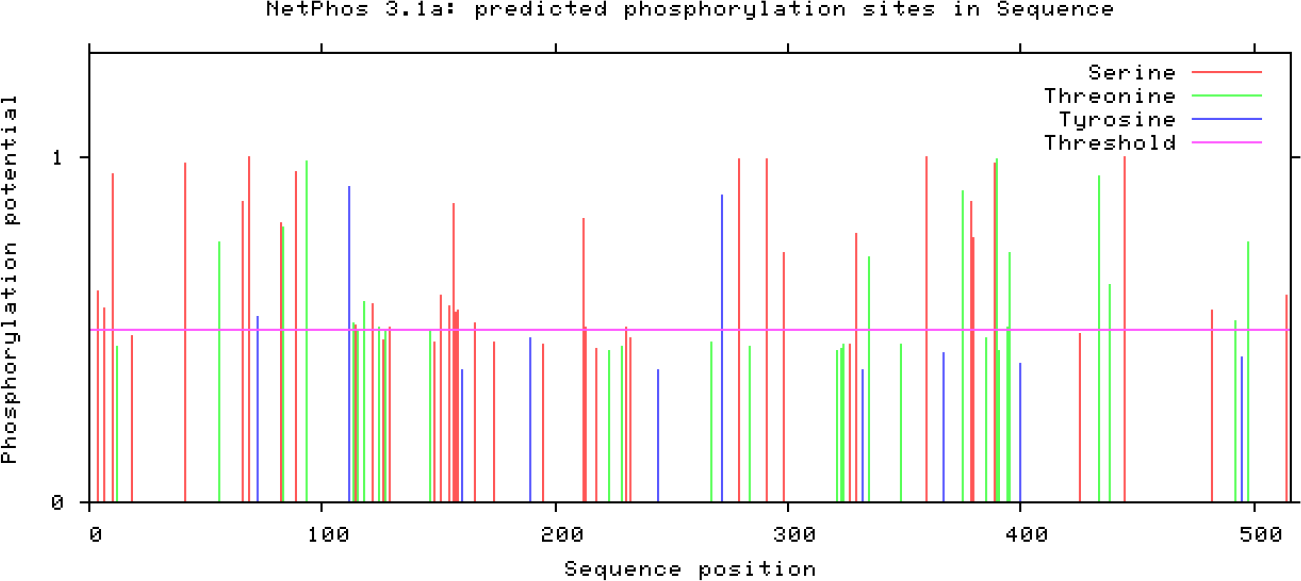
Tyrosine, serine & threonine phosphorylation sites in MATE1 protein

#### VII. Predictions for the effect of non-synonymous SNPs using SIFT

In this result we can visualize nonpolar amino acids as black, uncharged polar as green, basic as red and acidic as blue. There is no amino acid change and threshold values of all amino acids are above 0.05 which means there are no deleterious effects present in the protein (Figure 11).

**Figure 11:**
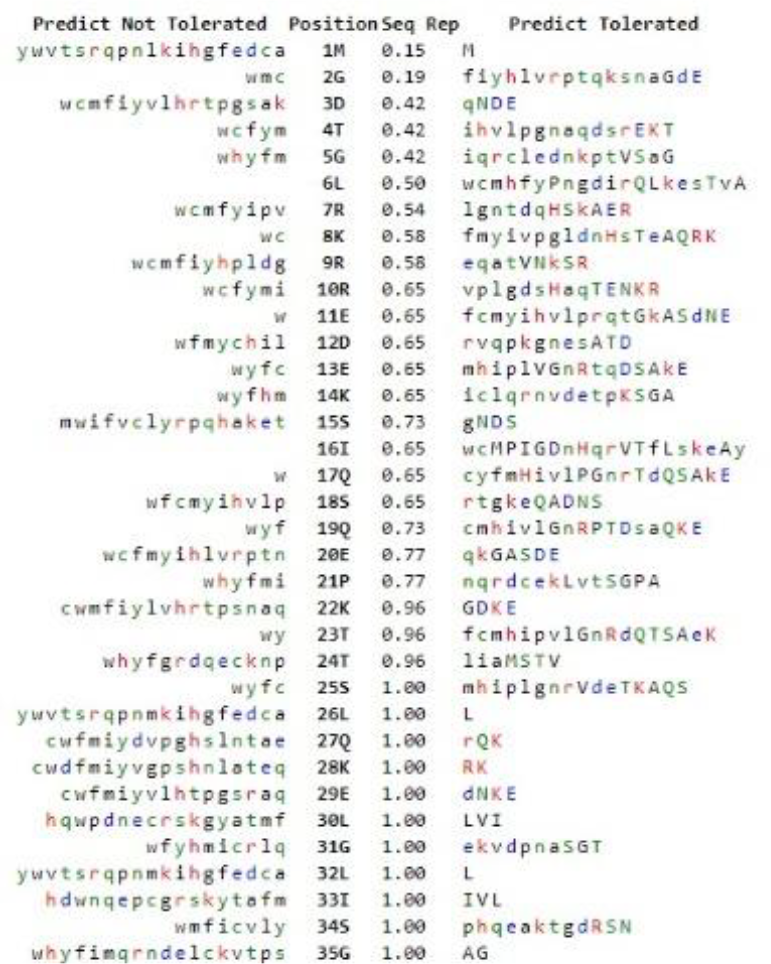
Non synonymous SNP predictions for CYP1A2 protein

#### VIII. Protein-protein interactions

Protein-protein interactions of CYP1A2 analyzed by STRING database showed protein and shell of interactions with other proteins such as CYP3A4, CYP2B6, CYP2E1, CYP2A6, CYP4A11, CYP7A1, EPHX1, GAPDH, NAT2, and UGT1A8 each represented with 11 different colored nodes in (fig 12). First shell is shown with colored nodes and the second shell with white nodes. The proteins with an unknown 3D structure are shown with empty nodes and the ones with known 3D structures are represented by filled nodes. Likewise, known interactions curated by the database are shown with blue edges and experimentally determined known interactions are shown with purple edges. The gene neighborhood is shown by green line, gene fusion with red lines, and gene co-occurrence with blue lines. The text mining interactions are shown with yellow edges, protein homology with sky blue and co-expression with black.

**Figure 12:**
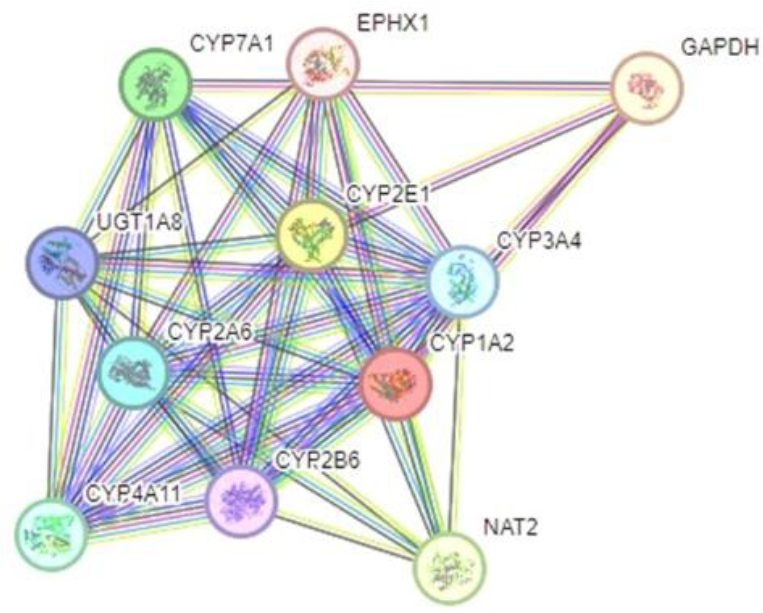
STING analysis of Protein-protein interaction of CYP1A2

Furthermore, statistics showed average node degree is 8.18 and PPI enrichment *p-value* is 7926e-12. Moreover, there are 45 edges and the local clustering coefficient is 0.896. There were 13 expected nodes and 45 different edges (Table 6).

**Table 6:** Protein-protein interactions of CYP1A2 protein.

|  |  |
| --- | --- |
| Number of nodes | 11 |
| Number of edges | 45 |
| Average node degree | 8.18 |
| Avg. local clustering coefficient | 0.896 |
| Expected number of edges | 13 |
| PPI enrichment *p-value* | 7926e-12 |

## Discussion

It is possible that CYP1A2 activity affects blood pressure by causing an increase in Na+ reabsorption in the proximal sector of the kidney’s nephron. [5]. CYP1A2 enzyme acts on the liver which induces caffeine metabolism resulting in caffeine metabolites (such as paraxanthine) which work in pair with caffeine itself to inhibit the adenosine receptors in kidneys as they both display antagonistic behavior towards the receptor. Adenosine is important for the regulation of water and sodium in the body and when it can no longer bind to the respective receptors, it results in decreased sodium and water retention in the body leading to low blood pressure. This process is illustrated below in (Figure 13)

**Figure 13:**
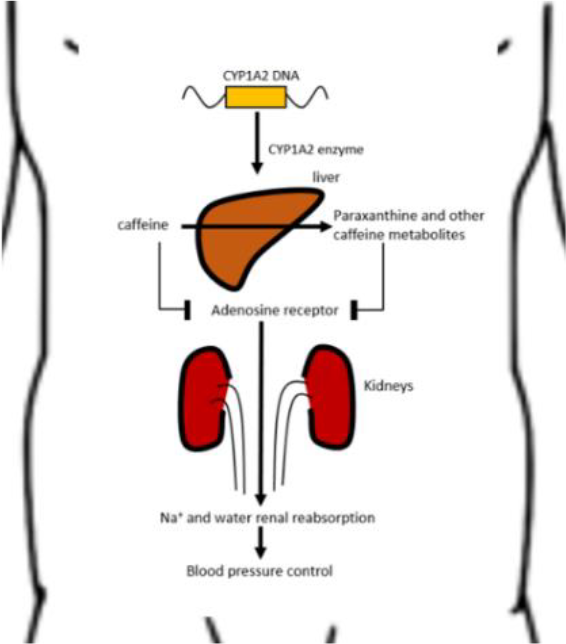
*CYP1A2* induced pathophysiology of caffeine metabolism

One Taiwanese study conducted by Hou Chien-Chou, et al. 2021, where 19,133 individuals were genotyped in which those with the same gene variant *CYP1A2* (rs762551), the subject variant showed a substantial association with coffee drinking. [9]. According to an Italian study, C allele variant of rs762551 demonstrated higher vulnerability to hypertension compared to A allele in Italian population[10]. A study conducted in Indonesia revealed that caffeine serves as the body’s primary source of antioxidants while simultaneously blocking adenosine A1 receptors. As a result, chemicals in foods high in caffeine that regulate pro- and anti-oxidants as well as inflammation levels may have a significant impact on blood pressure [11], which could be due to increase in heart rate, vasoconstriction or both. However, further studies must be conducted to make a sustainable conclusion. Moreover, individuals with C/C genotype, therefore slower caffeine metabolism, may have an increased risk of Myocardial Infraction (MI) [12].

KEGG (https://www.kegg.jp/pathway/map00232) pathway shows the caffeine metabolism pathway, names of different proteins and the metabolites involved. The contribution of CYP1A2 can be mapped by following the red highlighted boxes showing the protein’s involvement on every step. (fig 14)

**Figure 14:**
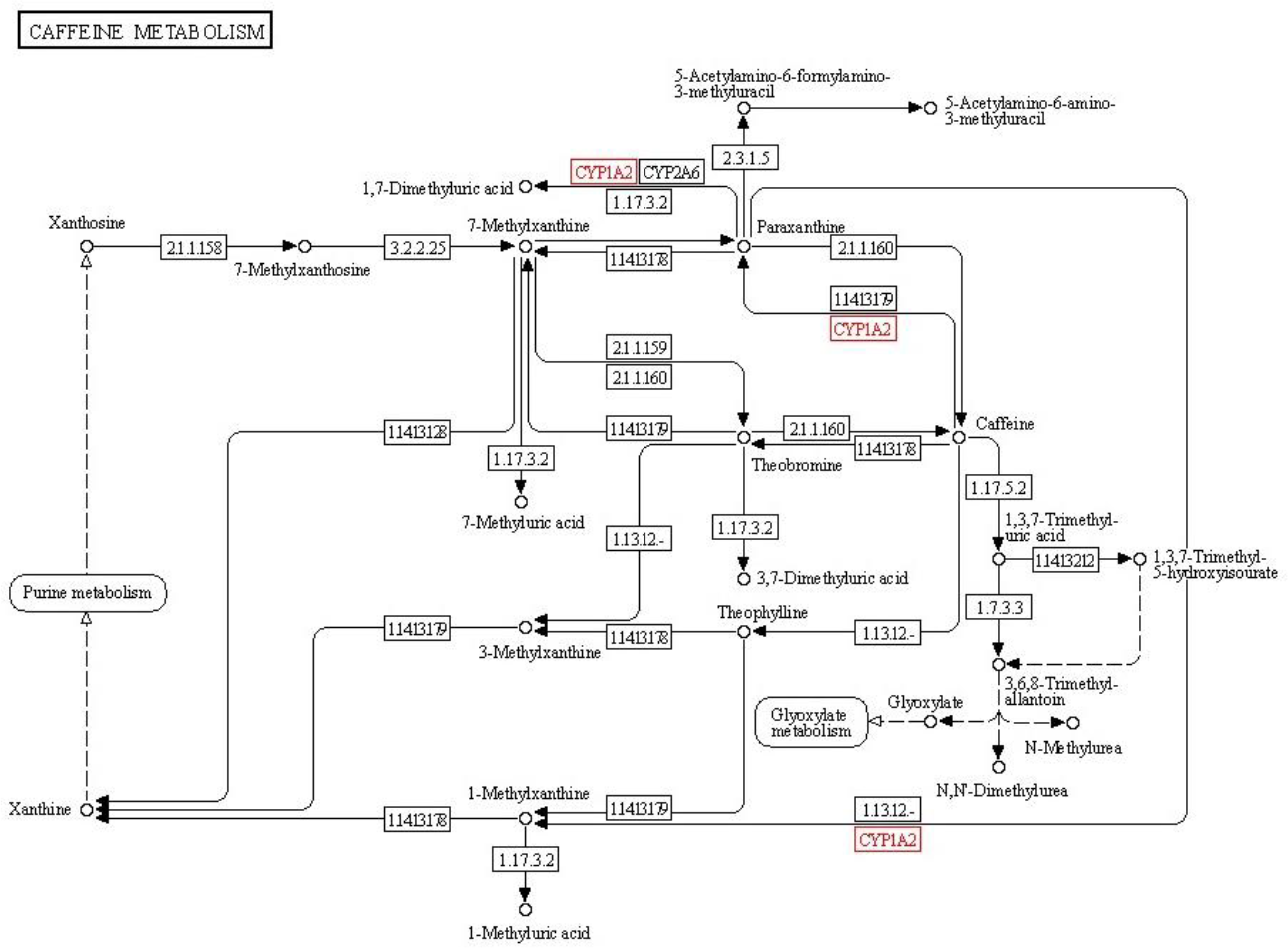
CYP1A2 protein involvement in caffeine metabolism

**Figure 15:**
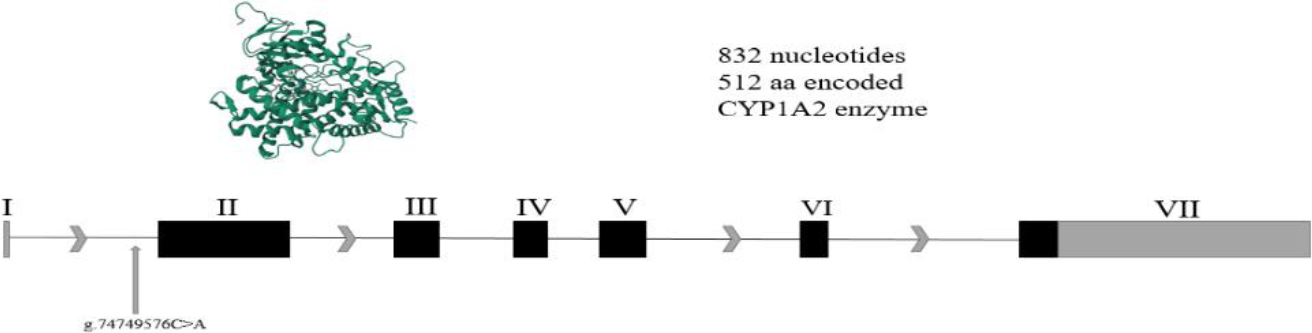
*CYP1A2* gene locus and crystal structure

CYP1A2 converts caffeine into Paraxanthine and further metabolizes it into 1,7-Dimethyluric acid. Paraxanthine is also converted into 1,3,7-Trimethyl-5-hydroxyisourate which is further metabolized by CYP1A2 into 1-Methylxanthine which is further processed into other products.

However, it is important to mention that *CYP1A2* is not the only gene responsible for producing a response against caffeine induced hypertension but it is the main enzyme responsible for caffeine metabolism. Moreover, factors other than *CYP1A2* polymorphism such as regularity of caffeine intake as well as physical fitness may play a role in hypertension [13].

## Conclusion

It is reported that the *CYP1A2* gene variant (rs762551) is found variable and associated with caffeine-induced hypertension (*p =* 0.039). However, other genetic factors/variants as well as the environmental factors may also play a substantial role.

## Acknowledgement

Authors are thankful for Al-Qaim Foundation, Gujranwala, Pakistan for providing the samples.

## Ethical Statement

The samples were collected by permission of Al-Qaim Foundation authorities by following the rules and regulations of the Institutional Review Board (IRB) of the hospital.

